# Hypothesis-free phenotype prediction within a genetics-first framework

**DOI:** 10.1101/2023.02.01.526358

**Authors:** Chang Lu, Jan Zaucha, Rihab Gam, Hai Fang, Ben Smithers, Matt E. Oates, Miguel Bernabe-Rubio, James Williams, Natalie Thurlby, Arun Prasad Pandurangan, Himani Tandon, Hashem Shihab, Raju Kalaivani, Minkyung Sung, Adam Sardar, Bastian Greshake Tzovoras, Davide Danovi, Julian Gough

**Affiliations:** MRC Laboratory of Molecular Biology; Bristol University; Shanghai Jiao Tong University; Centro de Biologia Molecular Severo Ochoa; King’s College London; Lawerence Berkeley National Laboratory

## Abstract

Cohort-wide sequencing studies have revealed that the largest category of variants is those deemed ‘rare’, even for the subset located in coding regions (99% of known coding variants are seen in less than 1% of the population^1–3^). Our understanding of how rare genetic variants influence disease and organism-level phenotypes has achieved limited progress, partly explained by the intrinsic difficulty in statistically evaluating the biological significance of rare events. Here we show that discoveries can instead be made through a knowledge-based approach using protein domains and ontologies (function and phenotype) that considers all coding variants regardless of allele frequency. We describe an *ab initio*, genetics-first method making molecular knowledge-based interpretations for exome-wide non-synonymous variants for phenotypes at the organism and cellular level. By using this reverse approach, we identify plausible novel genetic causes for developmental disorders that have eluded other established methods and present novel molecular hypotheses for the causal genetics of 40 phenotypes generated from a direct-to-consumer genotype cohort. This system offers a chance to extract further discovery from genetic data after standard tools have been applied.

## Main

Sequencing of human genomes holds great promise for using genetic information to guide medical discovery and therapy. And yet in general, advances in our ability to extract useful information from genetic data are not being made as rapidly as advances in our ability to generate the data, leading to a growing imbalance of effort. Systematically predicting potential organism-level phenotypes or disease risks based on the information of a person’s genetic variation remains an unsolved challenge. One important reason is the lack of consistent evaluation of consequences of genetic variants. The influence of common variants on phenotypes can be quantified by statistical weights from genome-wide association studies (GWAS) and presented as polygenic risk scores (PRS)^4–6^, while effects of rare variants can be expressed in terms of intolerance to high-penetrance deleterious variants in the human population^7–9^. Nevertheless, these two leave a large area in the effect size – allele frequency space under-explored (Fig. 1a), more explicitly not accounting for the influence of a significant amount of non-synonymous uncommon variants of medium or low penetrance^5^. On one hand, rare or low-frequency deleterious variants are explicitly removed by GWAS pipelines, and left without interpretation due to the inherent difficulties in statistical evaluation in population genetics. On the other hand, the statistical method for making gene-to-disease inference requires the support of multiple instances of high confidence predicted loss-of-function (pLoF) variants in a gene^7,10^, and it has been shown that such instances are rarely observed. It is estimated that cohorts roughly 1,000 times bigger than gnomAD (which contains 125,748 exomes) are needed to gather evidence of their existence in most genes^11^. On a few datasets with extensive broad phenotyping, PheWAS^12^ can leverage gene-based collapsing to address some rare variants. To further overcome these difficulties, and to apply to most datasets, there is demand for a non-association-based method; ideally one incorporating the effects of rare or low-frequency variants, expanding our capability for discovery within the constraints of the existing scale of human genetics data.

**Fig. 1.**
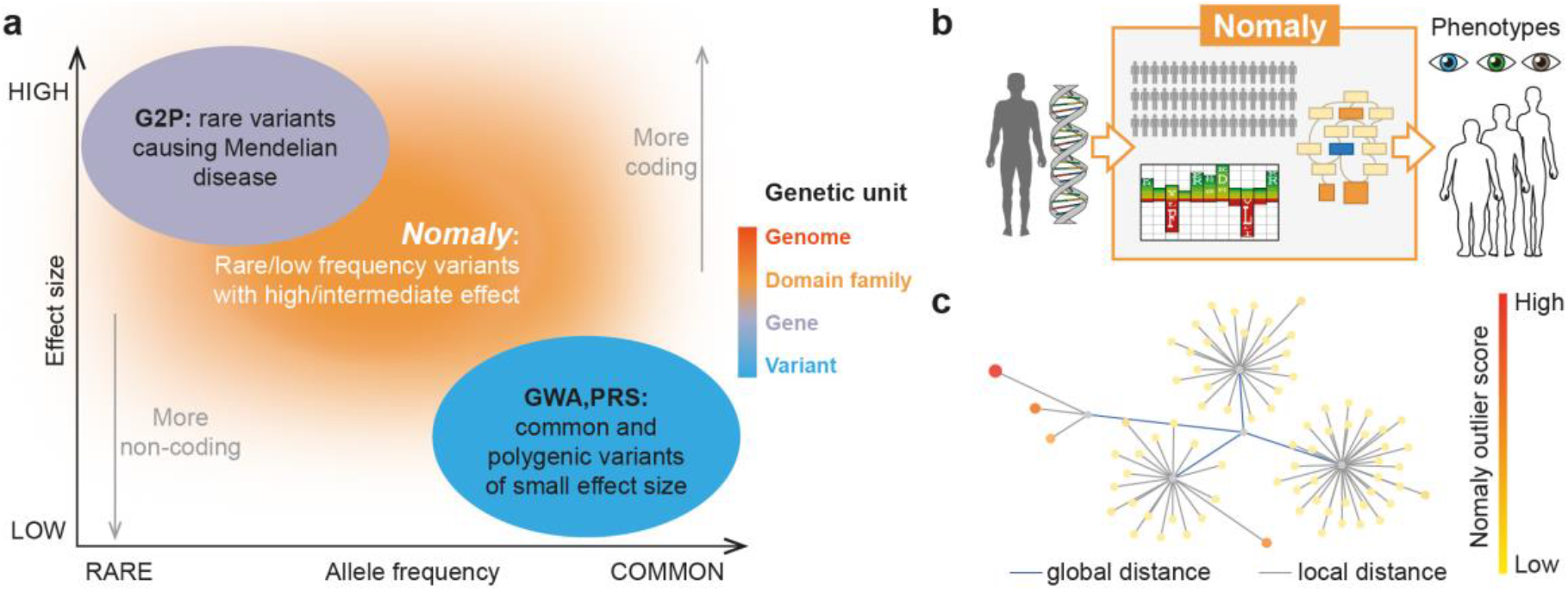
Framework incentive and design. (a) Positioning relative to heritability interpretation from prevailing genetic association analyses^5^. G2P: Gene-to-phenotype databases, GWA: Genome wide association, PRS: Polygenic risk scores. The colour bar shows the genetic unit of analysis employed by each method. (b) Framework overview. The method takes an individual’s genetic data as input and produces a list of ontology terms for which the person is a potential outlier. It uses a large background of genomes to which the individual is compared, ontology databases with gene-phenotype relationships, and evolutionary intolerance of mutations in protein domain families encoded by hidden Markov models (HMMs). (c) Schematic illustration of the genetic landscape for an ontology term (HP:0000834 ‘abnormality of the adrenal glands’) highlighting genomes with high outlier scores. Each node represents a genome, and edges are proportional to genetic distance in eigenspace – in essence, a reduced dimensional feature space between genomes.

Existing non-associative approaches that investigate genotype-to-phenotype relationships mainly use supervised network models, usually learned from large genomic variant databases focusing on specific phenotypes, e.g. antimicrobial resistance^13,14^, yeast cellular phenotypes^15,16^ and plant phenotypes^17^. These methods have demonstrated the possibility of using a knowledge-based strategy to make phenotypic prediction. However to achieve this, large datasets of millions of genetic variants (often through synthetic genetic arrays^18^) are needed and such datasets are only available for selected phenotypes, and such supervised models are not applicable to complex human phenotypes. We propose an unsupervised knowledge-based system (*Nomaly*), that makes *ab initio* predictions of potential phenotypes from thousands of ontology terms, leveraging knowledge of protein domains through hidden Markov models^19^. Instead of being used to train a model at an early step, the phenotypes are used as a final step to evaluate which predictions performed significantly better than expected by chance. The underlying protein knowledge on which the *ab initio* models are based can be examined to provide molecular insights into the predicted phenotype, in other words unlike supervised models, the interpretability of our predictions is high.

The *Nomaly* system (Fig. 2) is built on the premise that a genetic extreme outlier implies an outlier in phenotype. Under this hypothesis, the system evaluates the genetic heterogeneity in the context of each phenotype. Consequently, not only is it able to consider the additive effect of multiple variants but also the non-additive combinatorial effect where some variants become relatively rare and deleterious in the presence of other more common variants. The challenging computation of this is made tractable via a linear algebra approach solving an eigenproblem (spectral clustering), described as segmentation-based object categorization when used in image analysis^20^. A typical run includes a person or persons of interest and a large cohort-scale background, whereby outlier scores for thousands of terms in an ontology are calculated for each person of interest (Fig. 1b). The outlier scores represent the likelihood of being an extreme outlier in the ontology-specific genetic landscape with respect to the chosen background (Fig. 1c).

**Fig. 2.**
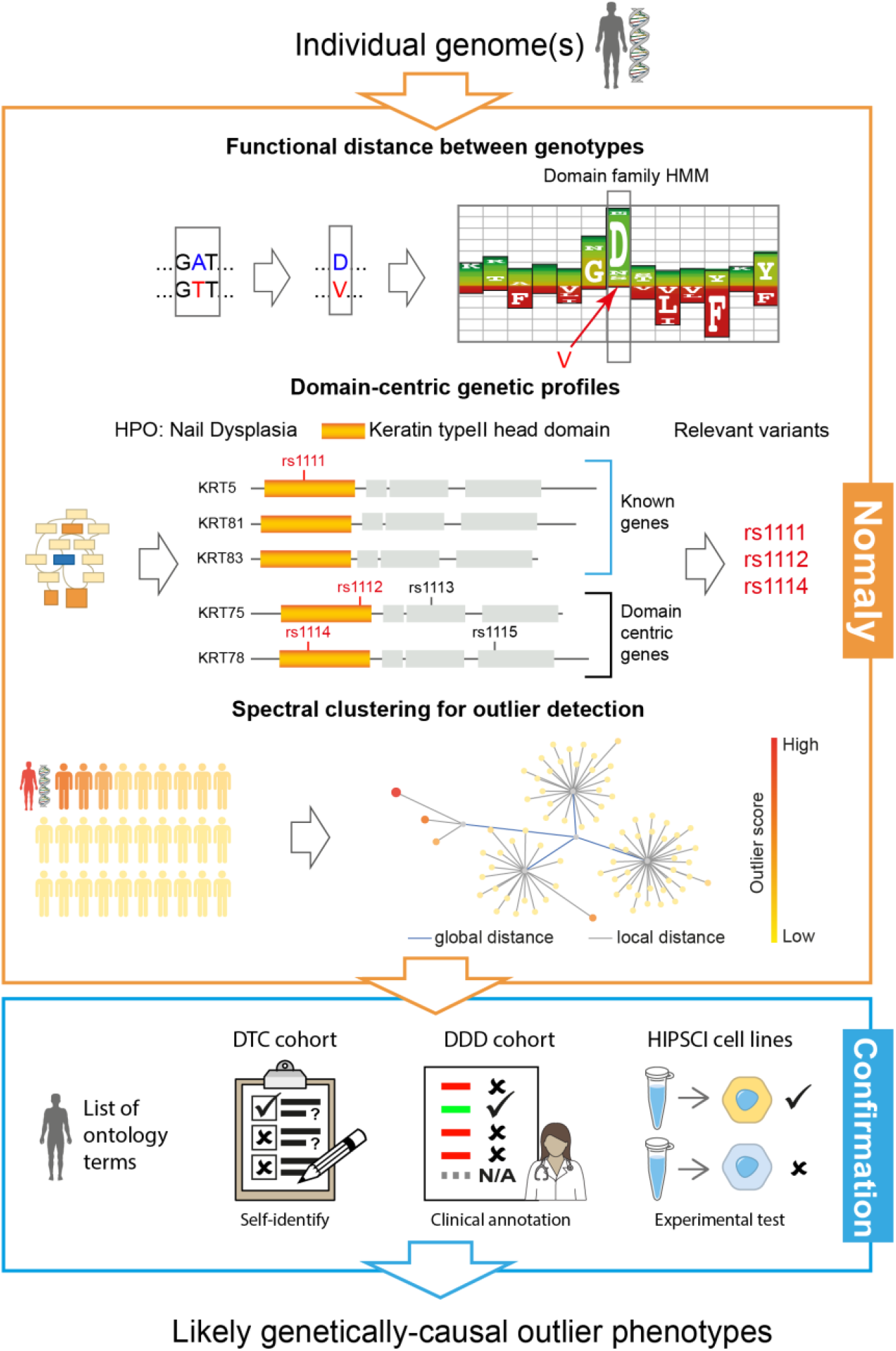
An outline of the analysis framework showing genome data as the input and likely genetically-causal outlier phenotypes as the output. In the orange top box (algorithm), firstly the functional distance between each missense variant is defined by the domain family HMM probabilities of the respective amino acids, scaled depending on zygosity. Subsequently domain-centric genetic profiles are constructed for each ontology term by collecting variants which are connected by common functional domain unit using dcGO^21^ enabling domain-based collapsing of a whole ontology term akin to gene-based collapsing used in PheWAS^12^. Lastly the profile defines the genetic distance to every genome in the background, and spectral clustering of the distance matrix identifies which genomes are outliers under the profile (HP:0000834); nodes represent genomes and are coloured by outlier score. In the next (blue) box, only the top-scoring outlier phenotypes are passed to the confirmation stage, where some of these genetics-first predictions are identified as likely plausible explanations of the recorded phenotype.

To evaluate performance, we wish to systematically assess: (i) whether there is a global consistency in statistics between the actual phenotypes and predictions based on outlier scores; (ii) whether the predictive success rate can be quantified; and (iii) how novel the predictions are. Below we demonstrate the significance and usefulness of our knowledge-based framework by answering these questions using three independent datasets, namely: a cohort of 2,248 participants specially recruited for this study (DTC, below), the well-established dataset of 1,133 children in the Deciphering Development Disorders (DDD) study with the respective gene-to-phenotype database DDG2P that provided genetic diagnoses for 40% of these children (majority through *de novo* mutations)^9,22,23^, and the Human Induced Pluripotent Stem Cells Initiative (HipSci) stem-cell bank^24^ where there is the possibility to experimentally verify predictions on cellular phenotype.

### Evaluation of the predictive power with a direct-to-consumer genetics cohort (DTC)

For the purpose of evaluation, we recruited a cohort of volunteers who had previously subscribed to direct-to-consumer (DTC) genotyping services (e.g. 23andMe, AncestryDNA and others), or who were otherwise already in possession of personal genomic data files to participate in this study (Extended Data Fig. 1). To test the overall significance of outlier scores we presented each participant with a questionnaire asking them to self-identify, from a set of questions, any phenotypes from among 25 of their top-scoring ontology terms mixed equally with a further 25 top-scoring terms from a decoy – a randomly selected individual from the background (Fig. 3a). The DTC cohort generated 2,248 questionnaires, yielding 94,966 yes/no answers across 3,672 ontology terms and 2,086 written comments. Questions were designed to identify only outliers, so if poor wording resulted in a common answer - observed as a positive response rate (>5%), they were excluded. By requiring participants to self-identify, the often-costly challenge of phenotype data collection is simplified, however we sacrifice accuracy, introducing noise that will mask the true predictive power by an unknown amount.

**Fig. 3.**
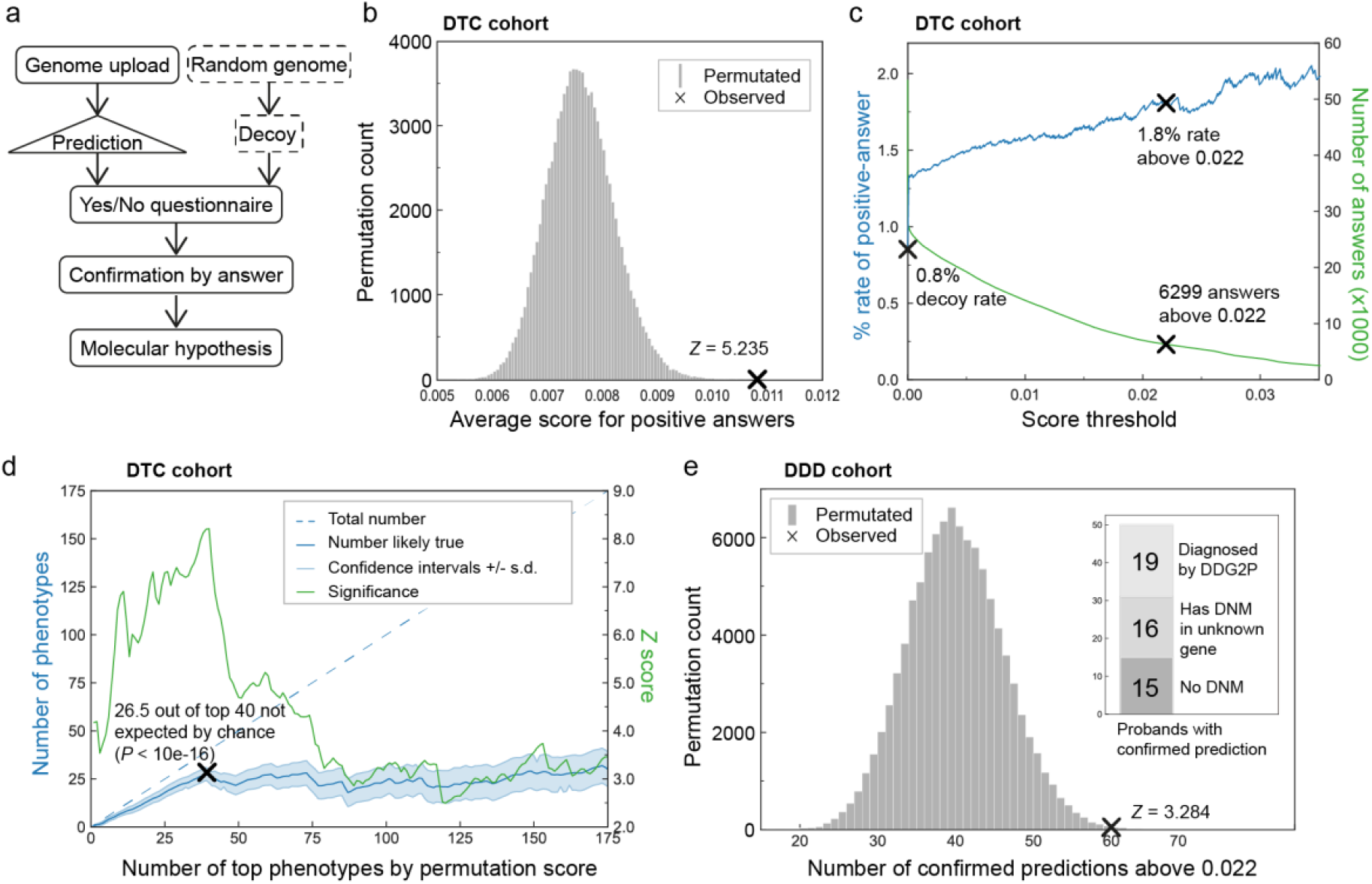
Evaluation of performance on DTC. (a-d) **and DDD** (e) **cohorts**. (a) Participants upload their DTC genome data on which outlier phenotypes are predicted, then shuffled with outliers from a decoy genome randomly selected from the background, to create a uniquely personalized questionnaire. Answers are used to confirm true predictions against decoys. (b) A test of 100,000 random permutations of the dataset shows that observed scores are on average higher for confirmed phenotypes, with a *p*-value of 8.25e-8 against randomly permuted scores. (c) The rate of identifying confirmed phenotypes by score threshold (blue) and number of above threshold predictions (green); at the default threshold of 0.022 the rate is more than double the rate for decoys, with a *p*-value of 7.93e-7 against 100,000 permutations. (d) The significance (green) of the top phenotypes by within-phenotype permutation of answers 100,000 times, and (blue) for the top *x* phenotypes, the number left after subtracting from the total those expected by chance (e) For DDD patients, the 60 above-threshold predictions confirmed by clinical annotation with a *p*-value of 5.12e-4 versus data from 100,000 random permutations. *Inset:* the 50 patients with top predictions compared to published data^22^ for whether a genetic diagnosis has been identified through DDG2P.

A statistical permutation test of outlier scores versus positive self-identification by participants proves that the method is significantly predictive of phenotype at the <1% level (*p*-value: 8.25e-8, Fig. 3b). The average rate at which participants self-identify a random phenotype was measured as 0.8% by taking answers to decoy questions. However, participants identify at a higher rate for predicted phenotypes and this increases monotonically with score threshold choice (Fig. 3c). For high-scoring phenotypes the rate of self-identification is 1.8% (*p*-value: 7.93e-7), but with improved phenotype measurement this could be increased. These results show that the high-scoring phenotype predictions harbour statistically significant signals and that about one half of the top-scoring predictions, verified through self-identification, are due to underlying genetic variation identified by the algorithm.

If a confirmed prediction has a genuine genetic basis and has not occurred by chance, then other predictions for the same phenotype in other participants are more likely to be true. This non-independence can be exploited and measured by per phenotype permutation tests. Further permuting across all phenotypes corrects for multiple hypotheses. Of the 40 phenotypes with the top statistic by permutation (Extended Data Table 1), only 13.5 are expected to have occurred by chance (Fig. 3d), which is a statistically much stronger result (*p*-value: <10e-16) than when considering predictions independently as above. Examples of a novel gene, recovery of a known variant, a novel variant in a gene related to a known gene, and mechanistic explanations are shown in Extended Data Fig. 2a-e.

### Novel potential genetic diagnoses for Deciphering Developmental Disorders cohort children (DDD)

In addition to the evaluation on our own DTC cohort for significance, a comparison was made to state-of-the-art work in the field on the well-established DDD cohort for the purpose of assessing novelty. DDD consists of 1,133 trios with developmental disorders who have been exome-sequenced and annotated with Human Phenotype Ontology (HPO)^25^ terms by clinicians ^9,22,23^.

Predictions on DDD were restricted only to HPO terms relevant to developmental disorders and used for annotation by clinicians. Of these, 60 predictions above threshold matching clinical annotations for 50 patients were found (Fig. 3e). Comparison to the published list of clinical diagnoses^9,22^ shows that 62% (31) of these patients had received no genetic diagnosis, including 15 who have no *de novo* mutations (DNM); the published diagnoses in the DDD paper were made through DNM missense variants, inherited variants and rare chromosomal events in known genes using the developmental disorder gene-to-phenotype (DDG2P) database. Thus, plausible genetic explanations can be discovered for families not covered by the established G2P interpretation method (Supplementary Table 2).

A global analysis of the predictions made on DDD that match clinical annotations confirms significance of the method (*p*-value: 3.62e-4), although it is less than for the DTC cohort due to limited data, restricted to developmental disorder-specific HPO terms. Likewise, about ⅓ of the above-threshold predictions (*p*-value: 5.12e-4) are expected to be true instead of ½ in DTC. Examples of a novel variant in a known gene and of a combinatorial effect are shown in Extended Data Fig. 2f-g. We conclude that, not only are the predictions significant on an independent dataset, but also largely non-redundant to those made by existing state-of-the-art methods, thus advancing the field.

### Application to interpreting genetic variants for cellular phenotypes on a large panel induced-pluripotent stem cell lines (HipSci)

Although HPO is the most common ontology used to annotate human phenotypes, and that used by DDD^22^, the DTC cohort study also included several other mammalian and disease ontologies (Extended Data Fig. 1b) and the gene ontology (GO)^26^. Despite being the ontology richest in data, GO performed worse on DTC than the other ontologies (*p*-value: 2.15e-2 for GO and *p*-value: 6.05e-7 for non-GO using threshold). This is presumably due to the difficulty in self-identifying molecular and cellular level terms, especially without recourse to invasive measurements on the person. The HipSci project provides exome sequence data for hundreds of iPS cell lines from different individuals, and thus offers an opportunity to make better use of the GO terms.

GO prediction was carried out on the exome sequences corresponding to 427 HipSci samples, generating predictions potentially relevant to cellular phenotype. It is not possible to use this dataset to assess global performance versus phenotype identification as with the other two cohorts, but the top-scoring phenotype terms were explored to identify a prediction that could be empirically validated *in vitro*. The phenotype “negative regulation of centrosome duplication” within the “biological process” domain of GO presented high-scoring predictions (Extended Data Fig. 2h) for some cell lines, and was selected principally for its suitability for measurement in iPS cells with an existing assay.

Centriole staining was carried out on HipSci cell lines corresponding to five individuals predicted to be outliers from their exome sequence, and three controls not predicted to be outliers. Counting of the centrioles indicated that three out of five predicted cell lines displayed an elevated percentage of cells with more than two centrioles, suggesting defects in centriole regulation and cell cycle, confirming the accuracy of the prediction (Fig. 4). This phenotype would not be identifiable from symptoms in the DTC or DDD cohorts, but this empirical evidence demonstrates the value of the approach for discovery at the cellular or tissue level.

**Fig. 4.**
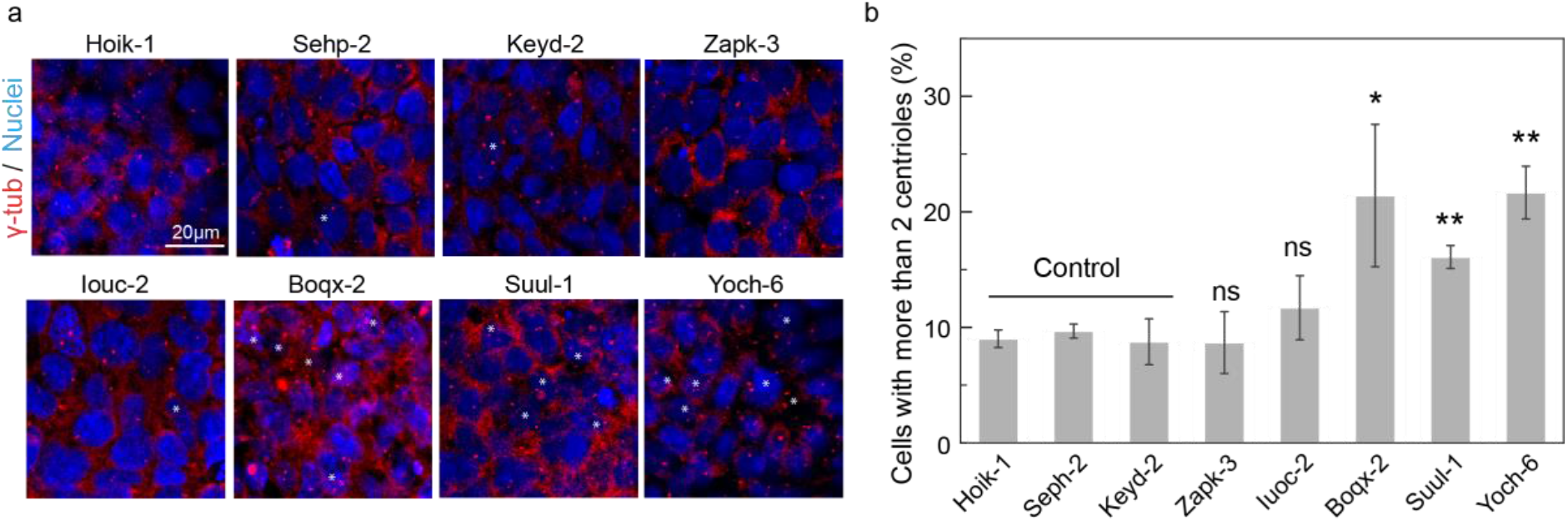
Testing for GO:0010826, which refers to ‘any process that decreases the frequency, rate or extent of centrosome duplication’. (a) Representative examples of each of the indicated cell lines. Centrioles were detected by staining with γ-tub. Asterisks indicate cells with more than two centrioles. The Hoik-1, Sehp-2 and Keyd-2 were control cell lines. (b) Histogram showing the percentage of cells with more than 2 centrioles per cell in the indicated cell lines. Results are summarised as the mean ± s.e.m. from 3 independent experiments (600-800 cells per cell line were analysed; *: P<0.05; **: P<0.005; ns: not significant).

### Types of genetic outlier that explain potential phenotype outlier

An outlier in the genetic landscape of an ontology term, as detected by this approach, can be caused by a single or by multiple rare variants. In the case of multiple variants, each variant can be classified according to whether it is absolutely required to achieve a score above threshold in the cohort, or whether it merely contributes to an above-threshold score; alternative variants can achieve above-threshold scores in different people (Fig. 5a,c). There can also be a combinatorial effect contributing to the score (e.g. Extended Data Fig. 2g), whereby a variant becomes relatively rarer and more deleterious in the presence of a particular genotype consisting typically of a few common variants, exemplified here by having a higher outlier score if classified into a cluster (Fig. 1c) than if there is no cluster. The most common case (71%; Fig. 5b, 1-a/b) is where a single variant that is highly deleterious is sufficient to achieve the threshold although others may contribute additional score (44% cases; 1-b). Multiple variants being required to cause an outlier account for 29% of cases (2-a,2-b). Combinatorial effects are important in a small minority of cases (Fig. 5d).

**Fig. 5.**
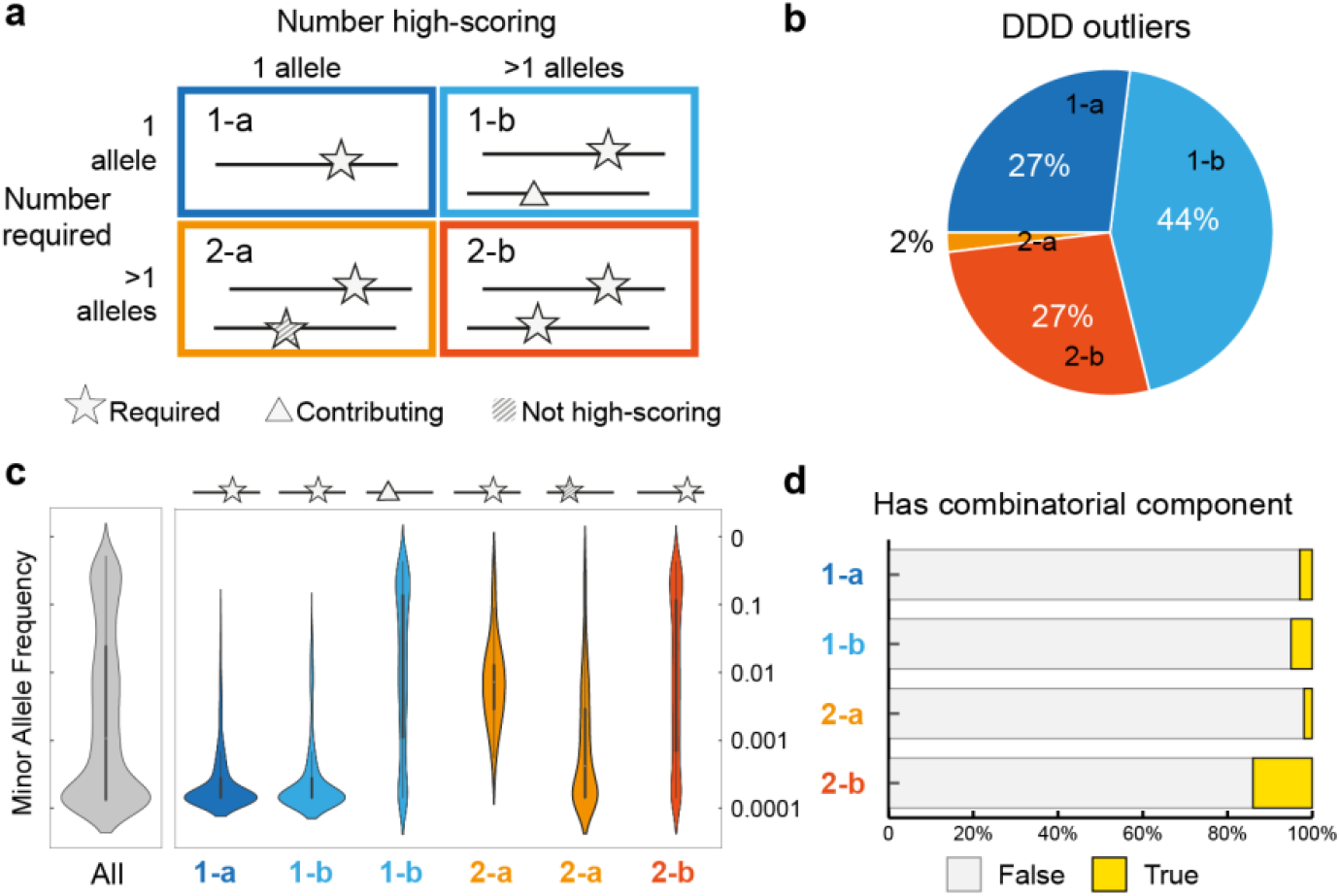
(a) Outliers classified by underlying genetic variants into 4 types: 1-a, single variant only required; 1-b, single variant plus contributing variant(s); 2-a, multiple variants but dominated by one high-scoring variant; or 2-b, multiple variants required. (b) Distribution of the 4 types of outliers in DDD. (c) The minor allele frequency of contributing variants for all outliers (grey), and by type (colours). (d) Percentage of outliers with a combinatorial component to the score with variants contributing non-independently.

The majority (90%) of single-variant-based predictions were caused by a rare variant with minor allele frequency (MAF) <0.5% for a heterozygous genotype, or MAF<1% for a homozygous genotype. However, not all rare variants result in a positive prediction, for example if it is not highly deleterious, or when many people from the chosen background harbour different rare variants for the phenotype, suggesting the ontology term is not highly evolutionary constrained. In multiple-variant-based positive predictions, 24% of variants are rare, and 55% are low-frequency (MAF<1% heterozygous or MAF<5% homozygous) (Fig. 5c). Whilst our approach does not filter variants based on allele frequency (as illustrated in Fig. 1a), there is insufficient power to recover the effects of common variants given the size of the DDD or DTC cohorts. As each phenotype question is only presented to a small subset of participants, only effects of large magnitude emerge from rare genotypes.

## Discussion

This paper describes a framework for performing and evaluating hypothesis-free phenotype prediction directly from a human genome. The key value is in providing potential genetic explanations for phenotypes that have been confirmed in the individual, which due to the high novelty and link to mechanism of the output, has potential for application to genetics-led drug target identification. Studying the combined effects on complex phenotype across variants in multiple genes is often impossible with simple statistical models, because of the lack of statistical power on existing cohort sizes. Also due to the lack of statistical power, low-frequency variants are usually explicitly excluded from GWAS. An *ab initio* model can partially overcome this limitation by testing a statistically small number of causally deduced direct predictions (a genetics-first approach). This knowledge-based framework with *in silico* and experimental validation approach described here has been shown to achieve this for exome missense variants. Our method, therefore, offers the ability to evaluate the effect of rarer genetic variants in a combinatorial way through linear algebra where statistical associative methods would not be applicable.

Hypothesis-free phenotype prediction with this genetics-first approach could be applied in principle to other *ab initio* models, but we chose to deploy a model based on protein domains, which are the functional units of proteins. Hidden Markov models built on protein domains^19^ enable the quantification of structural and functional effects of variants, and our domain-centric gene ontology (dcGO^21^) resource provides the link to phenotype. The domain-based model emphasises potential for novelty over coverage of known genes by carrying over the functional property of the domain across many genes.

Established genetic cohorts do not lend themselves well to testing a genetics-first approach since data is mostly only available for a restricted list of hypothesis-derived phenotypes – and usually not encoded in ontology terms, although increasingly attempts are being made to widen phenotype capture, e.g. using ICD-10^27^. The DDD cohort is one of the early large-scale studies to adopt HPO terms and was included in this analysis as a known reference point in the field. However, to truly test the genetics-first approach, a new DTC cohort needed to be created. Participants were recruited for their willingness to provide their genotype data first, then respond to personalised phenotype data collection post-prediction. Evaluation on both cohorts similarly confirms the predictions as highly significant yet also characterises the predictions as having a high false-positive rate; expectations are that for roughly ⅓ - ½ of confirmed high-scoring predictions, the causal genetic explanations will be true. Combining results per phenotype is more powerful, showing that explanations for about 26 of the top 40 confirmed phenotypes are likely to be true (Fig. 2d). The predictions were also found to be extremely high in novelty, confirmed by comparison to that achieved by DDD annotations of DNMs.

This method offers a powerful discovery tool for hypothesis generation from genetic data. The tool does not replace or compete with existing tools for human genetics which are largely aimed at being clinically actionable or offering effective intervention strategies. It rather complements and adds to them, aiming to enable medically high value scientific discoveries first suggested from the analysis of genomic data. Independent validation of the hypothesis generated by the prediction with relevant assays is recommended. The expectation is that results will be highly novel, characterised by high magnitude of effect from mostly rare variants, sometimes severally and occasionally acting combinatorially. In its essence the tool is a genetic outlier detector, so it is important to consider the background with respect to which the individual is an outlier. In this work we used the 1000 Genomes Project^1^ (1000G) genomes as a somewhat diverse background, but by selecting a different background, the definition of an outlier can be adjusted to suit the desired research question.

The catalogue of high-scoring results from the analysis of the three cohorts: DTC, DDD and HipSci provide over a hundred putative causal genetic explanations for numerous developmental, cellular and human/mouse/disease phenotypes. In addition to the extended data tables, they are also provided online with an interface to aid browsing and searching the variants, their classification of type, genes, scores and phenotypes (https:///supfam.org/nomaly), also including the database of 5,857 ontology questions as a resource for others.

The framework could also be used by other functional effect predictors, model organism ontologies and including known genes (to recover more of the less novel variants). Having proven principle on missense variants, we expect the future growth in this area by us and others will be by extension to other mutations e.g. indels and non-coding variants.

To sum up, the traditional approach to human genetics, where we ask “Does the data contain the answer to my question?”, has been turned on its head, and we instead ask: “For which questions does an answer lie within the data?”.

## Methods

### Nomaly framework

The framework consists of two primary parts: the predictive algorithm and the confirmation of predictions (Fig. 2). The algorithm takes an individual genetic data file (e.g. SNP array or exome) as input and outputs phenotypes for which the individual is predicted to be an outlier. The definition of ‘outlier’ is made relative to a background comprising thousands of genomes. For the DTC and HipSci cohorts the 1000G genomes were used as a background, but for individuals in the DDD cohort, the cohort itself serves as the background. There are thousands of potential phenotypes, taken from different ontology databases, each assigned a score for how much of an outlier it is for the individual in question against the background. In principle any genome or any biomedical ontology database can be used for a given study. Hidden Markov models (HMMs) are used to estimate deleteriousness; models from the domain databases SUPERFAMILY^19^ and Pfam^28^ were used, but in principle any HMMs can be used or any other measure of deleteriousness from a variant effect predictor.

The confirmation step is the assessment of whether an individual is indeed an outlier for a phenotype suggested by the genetics-first analysis. In principle the assessment can be carried out in any way that lends evidence to confirm the outlier (the DDD study used at least two certified clinical geneticists to perform assessment^22^) but in this work automated matching of ontology terms was used. Predictions from the first step that are subsequently confirmed in this step become candidate hypotheses linking a phenotype to variants via protein domains in genes.

#### Outlier scores

The predictive algorithm includes (1) quantification of consequence of missense variants using evolutionary intolerance to the amino acid substitution in a protein domain, done by taking the difference in amino acid emission probability (analogous concept to FATHMM^29^) and (2) generating variant lists for ontology terms through domain-centric linking to phenotypes (taken from dcGO)^19,21^. The mapping from variant to amino acids in proteins for (1) is done using the Variant Effect Predictor (VEP) tool^30^ (N.B. VEP used only for genomic mapping and no other functionality or scores are used) and mapping to domains using HMM sequence matching. The mapping of domains to ontology terms in (2) is combined with the variants falling within them from (1) to give a list of variants, commonly across multiple proteins, for each ontology term. Each ontology term is processed independently.

For a given term, a total genetic distance may be calculated between two individuals by summing the individual distances (defined as the log odds ratio of HMM probabilities for the two amino acids from the Dirichlet mixture) for all variants in the list for which they differ, depending on zygosity; distances are increased fourfold when both are homozygous and opposite. The all-against-all distances between the members of the background and the individual, and between each other, can be used to construct a distance matrix. This matrix is then translated to a similarity matrix through a Gaussian kernel and used as the input to spectral clustering to determine whether there is hidden structure in the genetic landscape of that term, namely by identifying the biggest gap in eigenvalues^20^. If no hidden structure is found, then the individual’s outlier score will be equivalent to the average Euclidean distance to members of the background; in this case individuals with very rare and highly deleterious genotypes will have a high outlier score (Fig. 5c). If a hidden structure is found by spectral clustering, K-means is then performed on the reduced-dimensional space derived from the top eigenvectors selected by the elbow method. In this scenario the outlier score becomes the sum of the local distance (from the cluster) and the global distance (between clusters) (Fig 1c). To normalise for cluster size the global distance is multiplied by μ where:

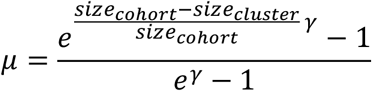

where *γ* specifies the penalty strength for large clusters; it was set to 9 in this study, conferring >99% penalty for large clusters with over 50% of the entire cohort.

Finally, since phenotypes have very different score distributions, a transformation (first described in Zaucha *et al*.^31^) is employed to generate a universal score function comparable between phenotype terms.

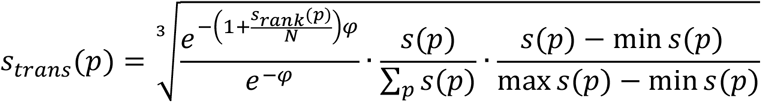

where *s_trans_*(*p*) is the transformed score of participant p in a given term, *s*(*p*) is the original score and *s_rank_*(*p*) is the rank of the original score within the Term. *φ* determines the strength of contribution of the rank to the total term score; it was set to 150 in this study, conferring the top ~2% a significant contribution.

#### Combinatorial component

The non-linear method of clustering used can result in one or more variants being markedly *rarer* within one cluster relative to the entire background, after genomes with other shared genotypes are grouped together. Thus a genome with a high global score due to the variants common to the cluster, and a high local score due to a variant rare only to that cluster, will have a higher total score than it would from its average Euclidian distance from the background. This additional score from variant rarity that only increases in the presence of other specific variants, is defined as the combinatorial contribution (as in Fig. 5d).

#### Computational cost

Calculating spectral clustering requires a large computation, even when using low level linear algebra BLAS^32^ libraries run on many threads in parallel. This is because it involves solving eigenproblems of a large matrix, which do not scale linearly with matrix size. For guidance, 5,800 terms on a batch of 50 SNP array genotype files with 2,504 background genomes (2554×2554 matrix) takes 21 hours with 12 threads (9.5 hours with 48 threads). Solving ~2,000 terms (independent eigenproblems) on a 3600×3600 matrix of exomes (including background) takes 3 hours using 600 threads.

### DTC cohort

#### Recruitment

Participants with access to personal direct-to-consumer genotype data were recruited anonymously online. Some participants were recruited in collaboration with OpenSNP^33^ and (separately) Sano Genetics. In this cohort 2,248 participations were recorded, with the participant website accessed from all over the world (Extended Data Fig. 1a). DTC genome data uploaded by participants was processed into a homogeneous format and quality-controlled with GenomePrep^34^, which also detects the sequencing method/genotyping array version. A similarity matrix of all genomes in the cohort was calculated, consanguineous relationships were recorded and genetic duplicates removed; people submitting independent files from multiple providers were only allowed to participate once (Extended Data Fig.s 1c-e). To prevent ‘gaming’ the study, for each participant a genome from the background was randomly selected, so the 25 ontology terms with the highest outlier scores for the participant could be randomly mixed with the 25 top-scoring ontology terms predicted for the background decoy genome to generate a personalised questionnaire with 50 questions. Each participant was invited to give a binary ‘yes’ or ‘no’ answer to whether they self-identify each phenotype, with the option to leave a comment (Extended Data Fig. 1b). In a small number of cases participants were recalled and invited to provide information supporting their answers for phenotypes of interest.

#### Process

The 1000G genomes were used as the background. Before recruitment, the binary yes/no questions were designed with the aim of identifying outlier phenotypes for 5,857 ontology terms, including terms from the Gene Ontology (GO)^26^, Human Phenotype Ontology (HPO)^25^, Disease Ontology (DO)^35^, Medical Subject Headings (MeSH)^36^, and Mammalian Phenotype (MP)^37^ ontology databases (Extended Data Table 1). See Supplementary Table 3 for the results or https://supfam.org/nomaly for an interactive version and database of ontology term to phenotype question mappings. Participant genome files were processed continuously in batches giving an approximately 4-12 hour turnaround between submission of file to generation of personalised questionnaire; results can be influenced by other genomes in the batch which effectively become part of the background for each other. This is because all parts of the similarity matrix interact with each other during solution of the eigenproblem.

#### Evaluation

We started with questions designed manually for 5,857 different ontology terms. In the end, 94966 binary self-identified answers were received for 3,672 questions, of which 1408 questions received at least one positive answer. Although questions were designed for participants only to self-identify when they are phenotypic outliers, often this was not achieved. To illustrate, a hypothetical bad question is “Do you have myopia?” whereas a good question would be “Do you have myopia worse than −6 dioptres”. Therefore, per-phenotype analysis was carried out for 342 ontology terms whose rate of participants answering ‘yes’ is non-zero and below 5%, and at least a total of 20 answers were recorded per term. Statistical evaluation was carried out using permutation tests with 100,000 iterations randomly re-allocating the outlier scores within the entire dataset or by ontology term.

### DDD cohort

#### Data source

Exome data from the DDD 1133 trio sequencing VCF files (accession code: EGAD00001001355), 1133 trio family relations, phenotype datasets, validated DNMs (EGAD00001001413), were obtained from the European Genome-phenome Archive at the European Bioinformatics Institute (EGA, https://ega-archive.org/).

#### Process

We ran the 1133 DDD probands, together with 1000G genomes as background, using HPO as the ontology database. Only variants that pass all filters (as specified in the meta-data information from EGA downloads) are included. HPO terms that were used to clinically describe phenotypes in the 1133 trio were mapped to HPO terms in the v1.2 database used in our predictions. Due to the nature of the method, running all of the probands at the same time, in one distance matrix, means that the background is a mixture of 1000G genomes and the other probands. If there was a common genetic cause shared by many probands, it would not get a high outlier score, but we assume enough genetic causes of the disorders are sufficiently independent to be eligible for detection.

#### Evaluation

For the DDD cohort, evaluation was automated through direct comparison, for each patient, of whether a high-scoring HPO term closely matches a respective clinical annotation. Defining a close match between ontology terms is challenging because distance over the ontology graph varies wildly in biological meaning; adjacent terms can be very similar or very different. To define closeness, a measure of information content through the graph is needed, for which we used the cumulative number of patients annotated by terms traversing the graph as a metric. The close matches between HPO terms are listed in Supplementary Table 4. For example we found the following 4 terms sufficiently similar to define as close to each other: HP:0004097 ‘Deviation of finger’ (2 probands), super-term HP:0009484 ‘Deviation of the hand or of fingers of the hand’ (1 proband), and sub-terms HP:0009179 and HP:0004209 – ‘deviation’ and ‘clinodactyly’ of the 5th finger (56 and 2 probands respectively).

It should be noted that the evaluation assumes all potential HPO terms in the DDD database are considered by clinicians, and that an HPO term is *not* true for the patient if it is not annotated. This represents an underestimation of true phenotypes, but is a necessary assumption for a fair and automatic evaluation.

#### Comparison to published diagnosis by DDG2P

For patients with true positive predictions, we checked against the list of diagnostic variants in DDG2P from the initial publication^9^ and the list of validated DNMs (obtained from EGA) to see if there is a genetic diagnosis and if there is a potential uninterpreted DNM. The 1133 trio DNM list showed that at least one DNM was found for 738 (65%) of children (excluding synonymous, intron, and intergenic DNMs).

### HipSci cohort

#### Data source

HipSci exome sequencing data for healthy and diseased people (EGAD00001003514, EGAD00001003521, EGAD00001003522, EGAD00001003524, EGAD00001003525, EGAD00001003516, EGAD00001003526, EGAD00001003527, EGAD00001003517, EGAD00001003161, EGAD00001003518, EGAD00001003519, EGAD00001003520) were obtained from EGA. HipSci open-access exome-sequencing data were downloaded directly from hosting website (https://www.hipsci.org/data).

#### Process

For each donor, one cell line was selected for the genetics-first prediction according to the following criteria: use primary tissue data if available, otherwise use the iPSC cell line with the minimum changes from the origin cell as measured by number of differences per Mbp, and excluding those where the pluritest or custom CNV check is missing. The result was that 437 cell lines from different donors were selected and the corresponding exome files processed using the 1000G genomes as background, predicting from a set of 5805 possible GO terms.

#### Evaluation

There was no confirmation step as with DTC and DDD, so no phenotype data was used. A list of phenotypes with several predicted outliers was examined to find one potentially verifiable by experiment. The list was initially narrowed to four candidates that could be tested on iPS-derived macrophage cells and two candidates that could be tested directly in iPS cells. Expression analysis of genes harbouring the variants of interest uncovered a lack of expression for GO:0002741 and GO:1900025 in macrophage, so these were eliminated. The variants implicated in terms GO:0035718 and GO:0002840 are involved in cell signaling (e.g. from the thyroid) and were eliminated as too experimentally complicated. One term, GO:1901223, was excluded due to lack of availability of a differentiated cell line for a key donor with the variant. Finally GO:0010826 was selected for experimental validation because five iPS cell lines predicted as outliers were available.

#### Experimental test for GO:0010826

The Hoik-1, Sehp-2 and Keyd-2 cell lines were selected as control. The Suul-1, Yoch-6, Boqx-2, Zapk-3 and Iuoc-2 cell lines were tested. Cells seeded at low confluence were grown for 2 days and stained for Ac-tub, γ-tub, and DAPI, with centrioles detected by staining with γ-tub. Results are summarised as the mean ± s.e.m. from three independent experiments (600-800 cells per cell line were analysed). Note that the cell lines Suul-1, Yoch-6 and Boqx-2 showed higher percentage of cells with more than two centrioles when compared to the control cell lines, indicating defects in centriole regulation and cell cycle. The cell lines Zapk-3 and Iuoc-2 did not show a centriole-related phenotype.

## Extended data figures and tables

**Extended Data Fig. 1.**
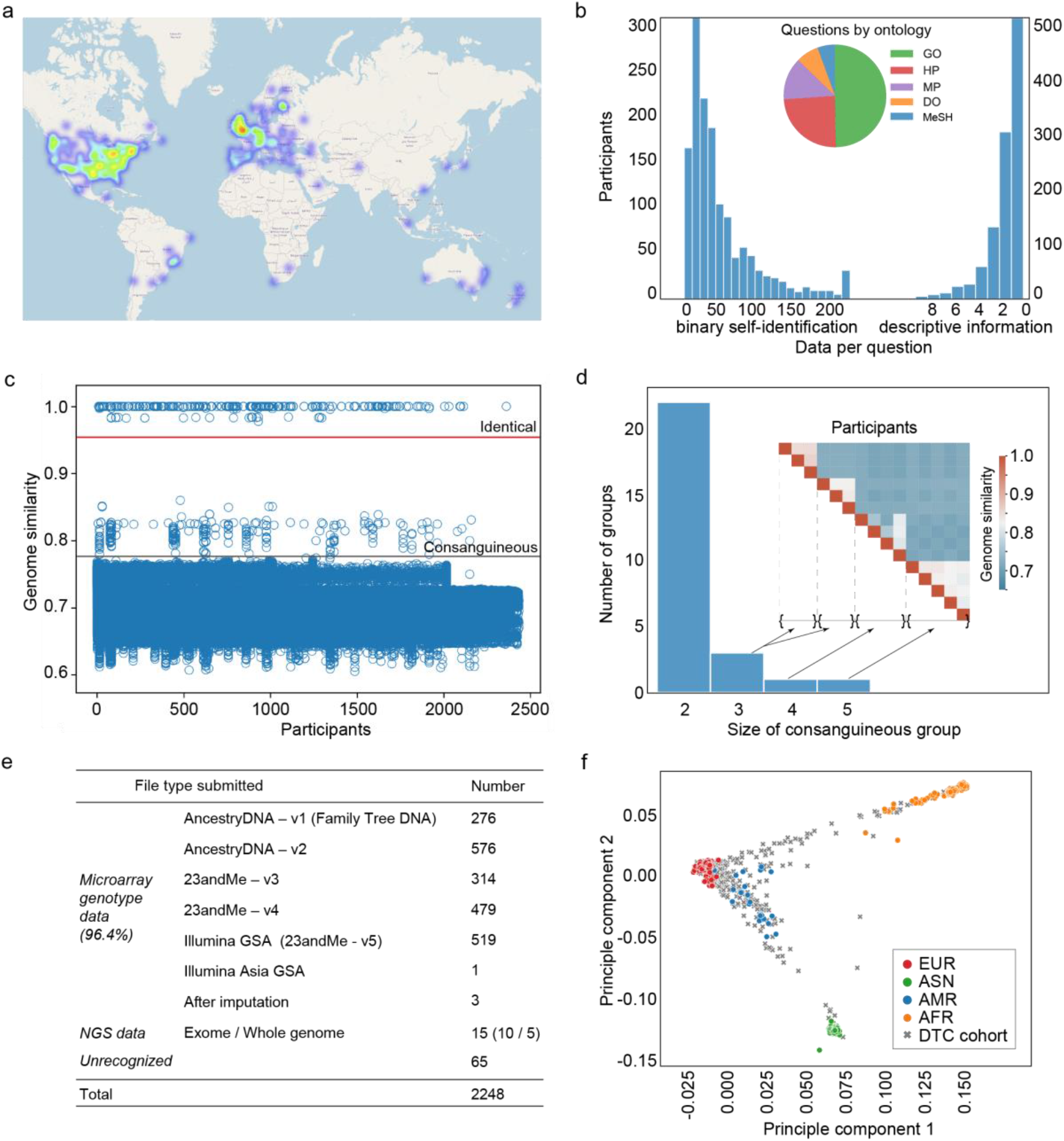
Overview of the direct-to-consumer (DTC) cohort. (a) The geospatial distribution of 2,248 participants. (b) Summary of the data collected for questions with at least one positive answer, including the number of binary answers to questions for self-identifying phenotype (left), number of descriptive free-text comments left per participant (right), and the breakdown of questions by ontology (pie chart) between: GO, gene ontology; HP, human phenotype ontology; MP, mammalian phenotype; DO, disease ontology; MeSH, mesh subject headings ontology. (c) Pairwise similarities between all DNA data files. Files are considered to be genetically identical if over 97% of variants are the same, and consanguineous if above 78%. (d) Consanguineous groups in the data. (e) The types and quantity of files submitted by participants, identified using GenomePrep^34^. (h) The ethnicity of participants mapped to the 1000 Genomes project ethnicity group principle components, labeled: EUR, European; ASN, East Asian; AMR, mixed American; AFR, African.

**Extended Data Figure 2.**
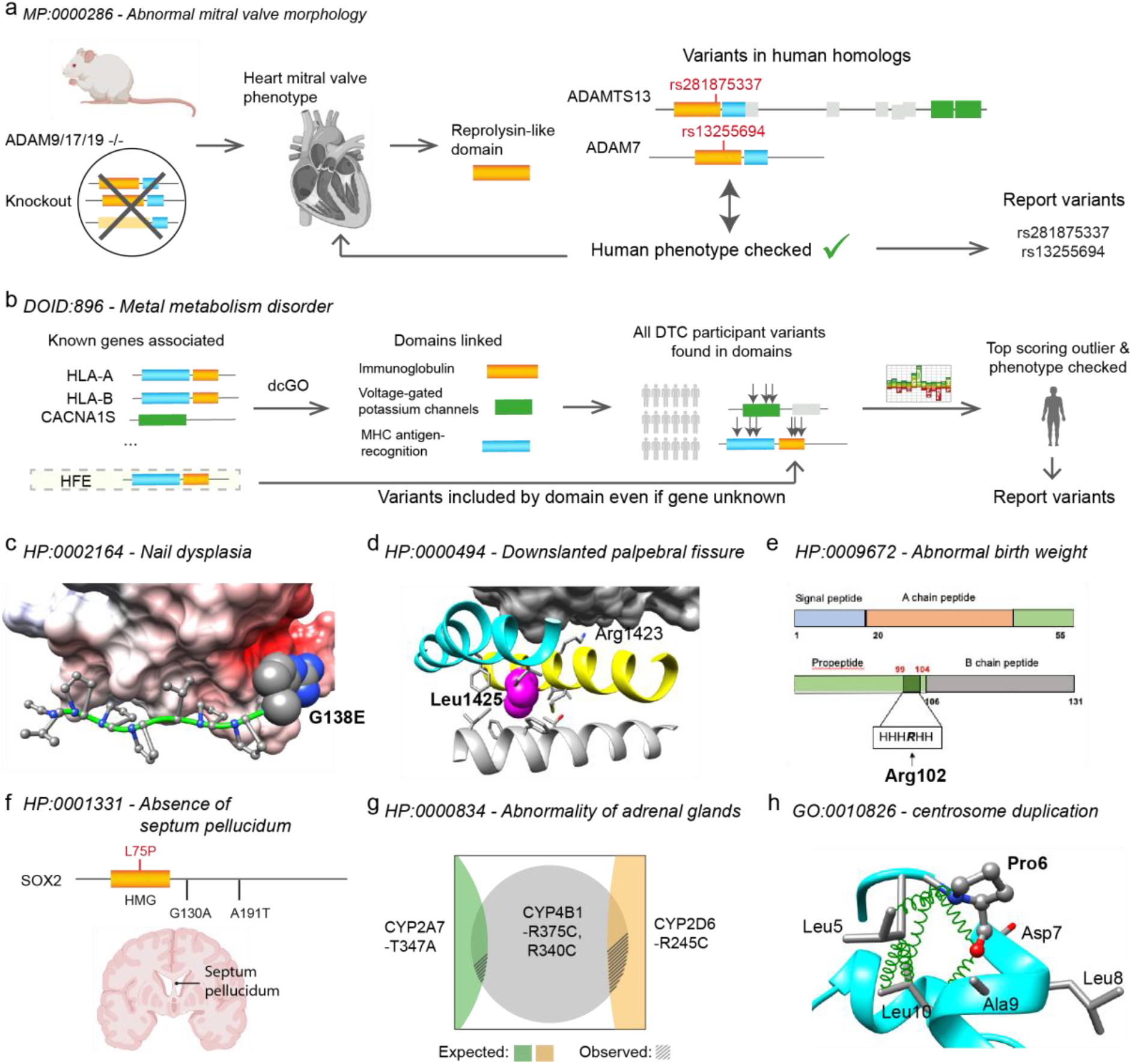
Examples. **(a)** *Novel gene association*. This representation shows how primary evidence from mouse knock-out experiments in genes ADAM9/17/19 led to a dcGO link between a mitral valve phenotype and the reprolysin-like domain shared by these genes. Novel variants identified in respective domain of human genes (ADAM7, ADAMTS13) confirmed the mitral valve phenotype via questionnaire. **(b)** *Known variant*. Chr6 Pos26093141 HFE-C282Y. A variant used to predict confirmed hemochromatosis in DTC participants was already known in ClinVar^38^. The diagram shows left to right: how multiple genes (including HFE) labelled with the ontology term were used by dcGO^20^ to link it to domain families, then variants in DTC participants within these domains prioritised by HMM probabilities led to prediction. Had the HFE variant not been known (HFE not included as known), it is possible with this approach that it would have been discovered *de novo* by the DTC study on the basis of domain association. **(c)** *Novel variant in related gene*. Chr12 Pos52913668, KRT5-G138E. The prediction includes other variants in keratin genes KRT75, KRT78 not previously linked to this ontology term. This variant is found in the head region (1-167) of the intermediate filament rod domain adjacent to an ELM motif involved in a protein-protein interaction with an SH3 domain. This figure shows a structural interpretation of the the mutation of G138 to a polar negatively charged Glu disrupting binding to the electrostatic surface of SH3 domain from PDB structure 2GBQ. **(d)** *Single variant*. Chr17 Pos29586054 NF1-L1425R. This single rare and deleterious variant in the neurofibromin (NF1) gene is sufficient to produce the top-ranked score for this phenotype in DTC. The image shows a structural interpretation of the central domain of neurofibromin (with helices 6c and 7c in cyan and yellow respectively) bound to Ras (surface model, top) and the Leu residue at 1425 rendered as spheres. Substituting Arg for Leu at this position will disrupt the helical geometry and successful interaction with Ras. **(e)** *Single variant*. Chr19 Pos17927755 INSL3-R102C. This single rare and deleterious variant in the insulin-like 3 (INSL3) gene is sufficient to produce the top-ranked score for this phenotype in DTC. The primary structure shows the variant lying within a conserved polar charged patch. The PDB structure 2H8B shows the two disulphide bonds that stabilise the structure. A mutation of the Arg to Cys could disrupt the polar charged region of the protein and even interfere with the correct formation of disulphide bonds. **(f)** *Novel variant in known gene*. Chr3 Pos18143037, SOX2-L75P. Two independent cases in the literature of missense mutations in SOX2 (G130A and A191T) mean this is already a known gene for the phenotype in the OMIM^39^ database. The phenotype was correctly predicted by our work on a DDD patient due to a novel variant in the HMG-box domain of this known protein. **(g)** *Combinatorial effect*. This phenotype correctly predicted on a DDD patient requires 2 variants in CYP4B1 (in linkage disequilibrium), plus additional variants. There is a combinatorial component to the score which raises this patient to the top rank in DDD for this phenotype. This Venn diagram shows that common variants are observed to co-occur (shaded areas) with the CYP4B1 variants much less than expected if the variants were independent (intersection areas). The top-ranked patient has all four variants. **(h)** *Experimentally validated on HipSci*. Chr3 Pos48414274 FBXW12-P6L. This homology model for F-box/WD repeat containing protein 12 (FBXW12) using PDB structure 1NEX as template shows the proline at residue 6 that mutates to leucine in the variant, residing within the N-Proline box motif in N-terminal F-box domain. A proline at the N-cap position mediates hydrophobic interactions similarly to other N-cap residues Asn, Asp, Ser, and Thr. The mutation to leucine in this position impacts interactions within the motif, likely causing a change of specificity and/or affinity.

**Extended Data Table 1.** The top 40 DTC cohort phenotypes and corresponding questions presented to participants (out of 5,857 possible questions). The first column includes the term ID, name and in brackets the number of positive ‘yes’ answers out of the total number of questions answered by participants.

| Term ID - name (answers: yes/total) | Question |
| --- | --- |
| DO:DOID:896 - metal metabolism disorder (1/46) | Metal metabolism disorder is an inherited metabolic disorder that involves metabolic disturbances in the processing or distribution of dietary minerals. Anyone in your family been diagnosed with metal metabolism disorder? |
| MP:0000286 - abnormal mitral valve morphology (2/50) | Abnormal Mitral valve regurgitation is a backflow of blood caused by failure of the heart's mitral valve to close tightly. Have you been diagnosed with abnormal Mitral valve regurgitation through Echocardiography (ECG) or Electrocardiography (EKG) test? |
| MP:0010402 - ventricular septal defect (1/47) | Ventricular septum is the wall dividing the lower chambers of the heart. Have you undergone Echocardiography showing any defect in ventricular septum? |
| GO:1901213 - regulation of transcription from RNA polymerase II promoter involved in heart development (2/85) | Have you ever had magnetic resonance imaging (MRI) showing thickened heart muscle which may indicate cardiac hypertrophy (abnormal heart muscle enlargement) which is characterized by shortness of breath, general fatigue, fainting, palpitations? |
| GO:0014854 - response to inactivity (3/80) | Do you have a medical history of polio attack or have you ever undergone electromyography (EMG) showing signs of nerve damage or neuromuscular examination showing signs of residual weakness and atrophy of muscles? |
| GO:0031646 - positive regulation of neurological system process (2/66) | Have you or anyone in your family ever undergone blood test showing very high levels of immunoglobulin E (IgE) which may indicate job's syndrome (immune disorder), characterized by recurrent cold, unusual skin rashes, severe lung infections? |
| MP:0009445 - osteomalacia (2/96) | Osteomalacia is the softening of the bones caused by impaired bone metabolism primarily due to inadequate levels of available phosphate, calcium, and vitamin D, or because of resorption of calcium. Have you undergone X-ray analysis showing the presence of osteomalacia? |
| HP:0000280 - Coarse facial features (1/83) | Does your face seems to be coarse like to having large, bulging head, large lips and tongue and small, widely spaced malformed teeth etc? |
| HP:0002605 - Hepatic necrosis (2/48) | Does your blood test shows that you have a elevated levels of liver enzymes which is the main symptom of hepatic necrosis or Have you been diagnosed with hepatic necrosis? |
| GO:0048708 - astrocyte differentiation (1/60) | Has any child in your family ever been diagnosed with Alexander disease (disorder of the nervous system) which is characterized by enlarged brain and head size (megalocephaly), seizures, stiffness in arms/ legs (spasticity), intellectual disability, developmental delay? |
| DO:DOID:11030 - corneal edema (3/60) | Corneal edema is the swelling of the cornea following ocular surgery, trauma, infection, inflammation as well as a secondary result of various ocular diseases. Have you been diagnosed with corneal edema? |
| GO:2000516 - positive regulation of CD4-positive, alpha-beta T cell activation (2/49) | Have you ever had blood test showing low level of albumin which may indicate hypoalbuminemia or Do you suffer from edema (buildup of fluid) in your legs or face, skin that's rougher or drier than normal, not having much of an appetite ? |
| GO:0016447 - somatic recombination of immunoglobulin gene segments (1/60) | Have you or anyone in your family ever had blood test showing decreased levels of immunoglobulins which may indicate immunodeficiency-centromeric instability-facial anomalies syndrome (ICF syndrome) (rare immune disorder), characterized by increased distance between bodily parts (hypertelorism), skin fold of the upper eyelid, unusually large tongue? |
| MESH:D010182 - Pancreatic Diseases (2/77) | Have you been diagnosed with any pancreatic diseases like pancreatitis (pancreas inflammation), pancreatic cyst, cystic fibrosis etc? |
| GO:2000104 - negative regulation of DNA-dependent DNA replication (2/113) | Have you ever been diagnosed with magnetic resonance imaging (MRI), shows defects in eye movements which may indicate ophthalmoplegia manifested by stroke, infection, abnormal eye movements, muscle weakness? |
| GO:0036314 - response to sterol (1/33) | Have you ever underwent electrocardiogram showing irregular electrical activity of heart and been diagnosed with heart attack which is associated with symptoms like tightness or pain in the chest, neck, back or arms, lightheadedness, abnormal heartbeat, anxiety, shortness of breath and shoulder discomfort? |
| GO:0000724 - double-strand break repair via homologous recombination (1/28) | Have you been diagnosed with photokeratitis (cornea inflammation) which is associated with symptoms like teary eyes, eye pain, swollen eyelids, feeling of sand in the eye, hazy and decreased vision ? |
| MP:0009701 - abnormal birth body size (2/73) | Anyone in your family have any abnormal body size during birth? |
| GO:0030510 - regulation of BMP signaling pathway (1/30) | Fibrodysplasia ossificans progressiva (FOP) - abnormal development of bone in areas of the body where bone is not normally present. Have you or anyone in your family ever had blood test showing increased level of alkaline phosphatase which may indicate fibrodysplasia ossificans progressiva or Do you have symptoms like loss of mobility or difficulty in speaking and eating due to inability to fully open the mouth, episodes of muscle swelling and inflammation ? |
| HP:0000204 - Cleft upper lip (1/28) | Have you noticed any of your family members have openings or splits in the roof of the mouth and lip or Do anyone have cleft upper lip? |
| HP:0000277 - Abnormality of the mandible (1/36) | Have you ever undergone X-ray analysis showing any abnormalities in lower jaw bone like broad jaw? |
| HP:0000269 - Prominent occiput (1/33) | Do you have prominent occiput or prominent posterior skull? |
| GO:0008333 - endosome to lysosome transport (1/24) | Arthrogryposis-Renal dysfunction-Cholestasis (ARC) syndrome is a multisystem disorder, characterized by neurogenic arthrogryposis multiplex congenita, renal tubular dysfunction and neonatal cholestasis with low serum gamma-glutamyl transferase activity. Have you or anyone in your family been diagnosed with Arthrogryposis-Renal dysfunction-Cholestasis (ARC) syndrome? |
| MP:0009672 - abnormal birth weight (2/57) | Incase of being a parent, Is your child have any abnormal body weight during birth? |
| MP:0001575 - cyanosis (4/86) | Have you noticed any bluish or purplish discoloration in your skin without any apparent cause or Have you been diagnosed with cyanosis? |
| GO:0006032 - chitin catabolic process (1/24) | Have you ever been diagnosed with Cystic fibrosis (CF) (genetic disorder) through Sweat test and genetic testing or do you suffer from long-term issues include difficulty breathing and coughing up mucus? |
| GO:0008592 - regulation of Toll signaling pathway (1/111) | Have you or anyone in family been diagnosed with familial cold autoinflammatory syndrome (periodic fever syndrome) which can cause recurrent, intermittent episodes of fever and rash that primarily follow exposure to cold? |
| HP:0000069 - Abnormality of the ureter (6/199) | Have you ever had ultrasonography analysis showing any ureter abnormalities like uterine cancer? |
| GO:0004792 - thiosulfate sulfurtransferase activity (1/27) | Have you ever had blood test or endoscopy analysis showing ulcerative colitis (colon ulcer) which may causes abdominal pain, diarrhea, weight loss, fever, anemia? |
| GO:0006021 - inositol biosynthetic process (3/64) | Have you ever been diagnosed with inositol deficiency which is characterized by symptoms like Vision and/or eye abnormalities, alopecia (patchy hair loss), fatty liver, loss of memory, constipation, high LDL cholesterol levels? |
| GO:0006284 - base-excision repair (2/47) | Have you ever had electromyography (EMG) or nerve conduction studies (NCS) or somatosensory evoked potential (SSEP) analysis resulted in nerve degeneration or Have you or anyone in your family been diagnosed with Huntington's disease (progressive breakdown of nerve cells in the brain)? |
| MP:0003920 - abnormal heart right ventricle morphology (2/65) | Have you ever had magnetic resonance imaging (MRI) scan that showing any morphological abnormality in heart right ventricle such as right ventricle degeneration, right ventricle aneurysm (weakening)? |
| HP:0000971 - Abnormality of the sweat gland (4/86) | Have you been diagnosed with any sweat gland disease like hyperhidrosis (excessive sweating), anhidrosis (inability to sweat normally)? |
| HP:0002921 - Abnormality of the cerebrospinal fluid (1/154) | Cerebral Spinal Fluid (CSF) analysis requires CSF which is collected by lumbar puncture. It is usually performed by a doctor who is specially trained to collect CSF. Have you had Cerebral Spinal Fluid (CSF) analysis showing any abnormality of cerebral spinal fluid like Hydrocephalus (accumulation of cerebrospinal fluid)? |
| GO:0032782 - bile acid secretion (1/52) | Intrahepatic cholestasis: impairs the release of a digestive fluid called bile from liver cells. As a result, bile builds up in the liver, impairing liver function. If you're pregnant women, have you been diagnosed with Intrahepatic cholestasis? |
| MP:0002896 - abnormal bone mineralization (3/62) | Have you had a bone mineral density test that showed abnormal deposition of bone mineral or Have you been diagnosed with bone mineralization disorders like osteomalacia (softening of the bones), osteogenesis Imperfecta (skeletal dysplasias)? |
| DO:DOID:2789 - parasitic protozoa infectious disease (1/49) | Have you ever underwent ova & parasite (O&P) test or stool test showing the presence of Giardia Lamblia (parasite) and been diagnosed with Giardia infection (parasitic infection) which is associated with symptoms like watery diarrhoea, greasy stools, fatigue, cramps, abdominal pain, bloating, malnutrition and indigestion? |
| GO:2000611 - positive regulation of thyroid hormone generation (3/143) | Have you ever had blood test and been diagnosed with Graves' disease which is characterized by anxiety and irritability, weight loss, despite normal eating habits, erectile dysfunction, bulging eyes, rapid or irregular heartbeat? |

## Supplementary information

Supplementary tables provide lists of plausible novel genetic causes from the study, and can be interactively searched in the online version (https://supfam.org/nomaly/).

**Supplementary–Table 1 - List of the top-performing ontology terms in DTC cohort.** (a) For each ontology term, showing the corresponding question, answer statistics, types of confirmed predictions, and contributing variants for predictions. The high-scoring variants were evaluated using combined information of rarity, zygosity and deleteriousness. Overall yes-rate below 5%. (b) Similar to a, control yes-rate below 5% (overall yes-rate above 5%).

**Supplementary-Table 2 - List of matched phentoype predictions in DDD cohort.** For each person with at least one clinically annotated HPO terms positively predicted, showing the matched annotation, the predictions that exactly/closed matched, types of matched predictions, and contributing variants. Diagnostic status: 0 if no DNM nor diagnosis is found for the person, 1 if DNM is found but not interpreted with DDG2P, 2 if a genetic diagnosis have been provided with DDG2P.

**Supplementary Table 3 (interactive version only) - Questions mapping.** Questions were specifically designed, such that people who are outliers of the phenotype described by corresponding ontology term tend to answer Yes. Ontology terms were selected from several ontology databases, including the disease ontology (DO), medical subject headings (MeSH), human phenotype ontology(HPO), mammalian phenotype (MP) and gene ontology (GO) databases.

**Supplementary Table 4 (interactive version only) - Mapping of HPO terms to closely matched ones used in DDD annotations.** Showing for each HPO term, the list of exactly or closedly matched term(s) that are used by the 1133 trio DDD phenotype annotations. Notice that over half of the HPO terms had no close match in DDD annotations (which is 30% of those included by Nomaly).

